# MgAl-layered double hydroxide nanoparticles mediated oral delivery of dsRNA silencing *jhp* and *chi* genes with sublethal effects in *Phthorimaea absoluta*

**DOI:** 10.1101/2025.10.17.683103

**Authors:** Ayaovi Agbessenou, Jiangbin Fan, Eva Hotz, Katja R. Richert-Pöggeler, Christian Borgemeister, Jehle Johannes A.

**Author notes:** Key Laboratory of National Forestry and Grassland Administration for Control of Forest Biological Disasters in Western China, College of Forestry, Northwest A&F University, Yangling, Shaanxi Province, China (Jiangbin Fan).

## Abstract

**BACKGROUND:** The use of RNA interference (RNAi)-based technology for the management of agricultural insect pests is gaining momentum worldwide. A prerequisite for successful control by RNAi is the formulation of a highly efficient and stable double-stranded RNA (dsRNA). Here, we investigated the use of magnesium aluminum-layered double hydroxide (MgAl-LDH or LDH) nanoparticles complexed with dsRNA, termed BioClay^TM^, to target the *juvenile hormone inducible protein* (*jhp*) and the *chitin synthase A* (*chi*) genes of the South American tomato pinworm *Phthorimaea (=Tuta) absoluta*, a key pest of tomato crop.

**RESULTS:** Degradation assays showed that higher amounts of RNase A degraded dsRNA complexed with the LDH nanoparticles while lower RNase A amounts did not degrade the dsRNA. Furthermore, our findings revealed that dsRNA is stable after 60 min of incubation when in buffer solutions at pH 5 and pH 9, while naked dsRNA showed slight degradation at pH 11. Also, BioClay^TM^ and naked dsRNAs targeting *jhp* and *chi* genes did not significantly increase larval mortality in *P. absoluta*, but induced sublethal phenotypes through reduced pupal weight and adult emergence. Gene expression analysis revealed significant reduction of 73% and 48% in transcript abundance of chi gene in larvae that fed on *chi*-dsRNA and BioClay^TM^ *chi*-dsRNA after 72 h, respectively, while a 39% reduction in transcript of jhp gene was recorded only in larvae fed on BioClay^TM^ *jhp-dsRNA*.

**CONCLUSION:** Our findings provide critical insights into the significant impact of the use of dsRNA technology for targeted gene disruptions affecting important traits in *P. absoluta* and highlight the potential and opportunities that remain to be explored in the formulation and successful delivery of dsRNA-based active substances in pest management.

**Graphical Abstract:** 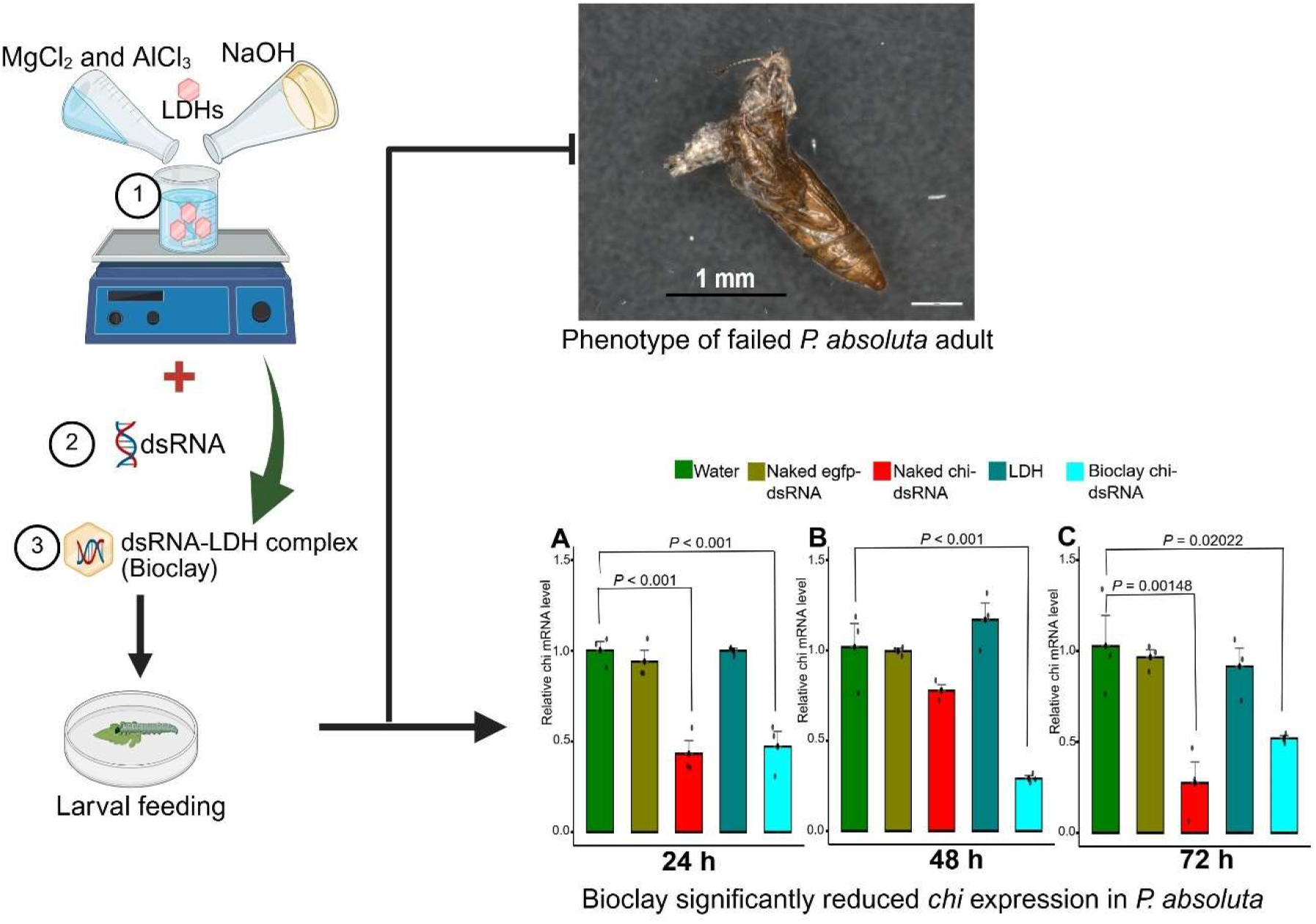

MgAl-LDH nanoparticles used as carrier to deliver dsRNA-based active substances hold potential to target *jhp* and *chi* candidate genes in *Phthorimaea absoluta* for the sustainable management of the pest.

## 1 INTRODUCTION

The South American tomato pinworm, *Phthorimaea* (=*Tuta*) *absoluta* (Meyrick) (Lep.: Gelechiidae) is a native insect pest to South America which was first reported outside its region of origin in eastern Spain in 2006, where it causes substantial yield losses to solanaceous crops.^1,2^ The pest larvae are the most damaging livestage and are known to voraciously mine the leaves, leading to significant yield losses of up to 100% if left uncontrolled.^3,4^ Currently, synthetic insecticides remain the most widely used control method against the pest, which have, however, serious limitations due to their negative impact on human and animal health, and the environment in general.^3^ Additionally, overreliance on insecticides has led to the development of resistance in *P. absoluta* populations.^5,6^ Therefore, the need to develop and deploy biological control methods, compatible with other more eco-friendly pest management practices.^7–10^

As a new technology, the use of RNA interference (RNAi) is gaining increasing attention worldwide as a novel pest control strategy.^11,12^ RNAi is a widespread post-transcriptional silencing mechanism of messenger RNA (mRNA), occurring in eukaryotic organisms where the triggering molecule, double-stranded RNA (dsRNA), regulates gene expression by targeting specific endogenous mRNA molecules in a sequence-specific manner.^13^ Upon cellular uptake, dsRNA molecules activate the RNAi pathway, a natural antiviral defense mechanism that processes long dsRNAs into small interference RNAs (siRNAs).^14^ These siRNAs are guided by RISC (RNA-induced silencing complex) to their complementary sequence in mRNA which is cleaved and prevented from translation, resulting in a reduction in gene product; inducing the gene silencing. The success of RNAi-based pest control strategy depends on the ability of insects to orally ingest the pest specific-dsRNA from the environment; hence, the dsRNA should be efficiently taken up by insect cells to access the RNAi machinery.^15^

Attempts to induce RNAi in *P. absoluta* by oral delivery of dsRNA suggested that RNAi holds considerable promise through the knockdown of essential genes responsible for the growth and development of the pest.^16–18^ For example, studies have targeted genes such as *Vacuolar ATPase-A* (v-ATPase-A), juvenile hormone-related proteins, and chitin synthase, employing diverse delivery methods like injection, root absorption, and topical applications.^19,20^ However, the efficiency of RNAi in lepidopteran insects, and especially in *P. absoluta*, is extremely variable and dependent on several factors including the alkaline nature of the midgut, target gene (sequence and length), mode of dsRNA delivery and stability of dsRNA to extracellular degradation.^21^ These factors make the selection of a suitable delivery method important for successful RNAi-based pest control. Consequently, several innovative strategies are being developed to address the delivery challenges and include nanoparticle-based systems which serve as dsRNA carriers.

Layered-double hydroxide nanoparticles (LDH), commonly known as clay nanosheets, have emerged as versatile carriers for delivering dsRNA.^22,23^ They possess favorable physical properties with low cytotoxicity, and serve as highly efficient delivery vehicles for nucleic acids. For example, Jain *et al*.^22^ reported the use of LDH nanoparticles as suitable carrier for the delivery of dsRNA-based active substances against the whitefly, *Bemisia tabaci* (Gennadius) (Hem.: Aleyrodidae) through foliar application, representing an attractive avenue for the sustainable management of the pest. Similarly, Lichtenberg *et al*.^24^ tested the ability of different nanoparticles to deliver dsRNA and observed that only Mg−Al layered double-hydroxide nanoparticles were effective at gene knockdown in *C. elegans*. Our study aimed at exploring the potential use of LDH nanoparticles as carrier to deliver dsRNA to *P. absoluta* targeting the *juvenile hormone inducible protein* (*jhp*) and the *chitin synthase A* (*chi*) genes. Specifically, we assessed the dsRNA degradation activity, then tested the RNAi efficiency against the candidate target genes.

## 2 MATERIALS AND METHODS

### 2.1 *Phthorimaea absoluta* rearing

A *P. absoluta* colony was set up at the Institute for Biological Control, Julius Kühn Institute, Dossenheim, in July 2023 using eggs and pupae kindly provided by Andermatt Biocontrol (Switzerland). Insects were kept on tomato plants (*Solanum lycopersicum* cv. Red Robin) as previously described by Agbessenou *et al*.^25^ Briefly, *P. absoluta* adults were maintained in ventilated cages containing tomato plants in a rearing room at 24 °C constant temperature with 50% relative humidity and a photoperiod of 16:8 h (light: dark). Adult moths were kept in ventilated cages (60 × 60 × 90 cm) and were fed with 10% honey solution placed on the top side of each cage.

### 2.2 *Phthorimaea absoluta* RNA extraction and cDNA synthesis

To clone the target genes, *chi* and *jhp*, total RNA was extracted from third and fourth *P. absoluta* instar larvae using the NEB Monarch Spin RNA extraction kit (NEB, T2110) following the manufacturer’s instructions. Then, a NanoDrop 2000c spectrophotometer (Thermo Fisher Scientific Inc., Waltham, MA, USA) was used to assess the quality and purity of the extracted RNA. cDNA was synthesized with up to 1,000 ng of purified RNA in 20 µL reactions using the First Strand cDNA Synthesis Kit (Thermo Fisher Scientific, Waltham, MA, USA).

### 2.3 Selection, amplification, and cloning of candidate genes for silencing in *P. absoluta*

Based on previous studies exploring the use of RNAi technology in *P. absoluta*, two candidate genes were selected including *jhp* and *chi*.^19^ Target regions were selected considering gene specificity, essentiality, size and region to obtain dsRNA. Fragments of 168 bp and 419 bp of *P. absoluta* target genes *jhp* and *chi*, respectively, were amplified from larval cDNA using specific primer pairs as described by Camargo *et al*.^17^ Amplification products were analyzed on 1% agarose gel electrophoresis in TAE buffer. Bands with expected molecular weight were cut and purified using the Zymoclean™ Gel DNA Recovery Kit (Zymo Research, Freiburg, Germany). The fragments were quantified and cloned into pGEM-T Easy vector (Promega, Madison, WI, USA) according to the manufacturer’s instructions, generating pGEM-*jhp* and pGEM-*chi*. The ligation products were transformed into competent *Eschericha coli* DH5α cells by electroporation. The insertion fragments were Sanger-sequenced and aligned against the *jhp* and *chi* sequences, in order to confirm no mutation during gene cloning. The fragment of 500 bp of the *egfp* gene was cloned into pGEM-T vector as control, generating pGEM-*egfp*.

### 2.4 Double-stranded RNA synthesis

pGEM-*jhp*, pGEM-*chi*, and pGEM-*egfp* genes were used as a template for transcription *in vitro* to produce dsRNA using gene-specific primers incorporating a T7 promoter sequence (TAATACGACTCACTATAGGG) at the 5’ end (Table 1). PCR conditions were 95 °C for 1 min, then 35 cycles of 95 °C for 15 s, 55 °C to 62 °C for 15 s, 72 °C for 10 s, and an extension step at 72 °C for 10 min. Amplified fragments were run on and purified from 1% agarose gels as described above. dsRNA was synthesized using a T7 RiboMAX Express RNAi System Kit (Promega, Cat# P1700) according to the manufacturer’s instructions, using 1 µg of gel-purified T7 PCR product as a template in a 20 µL reaction. The reaction mixture was incubated at 37 °C for 4 h and heated at 70 °C for 10 min, and double strands were allowed to anneal at room temperature for 20 min. Subsequently, dsRNA was treated with RNase and DNase I to remove the templates and purified with 95% ethanol. The purified dsRNA was quantified with a NanoDrop 2000c spectrophotometer (Thermo Fisher Scientific Inc., Waltham, MA, USA), and the integrity of dsRNA was subsequently screened on a 1% agarose gel.

**Table 1.**
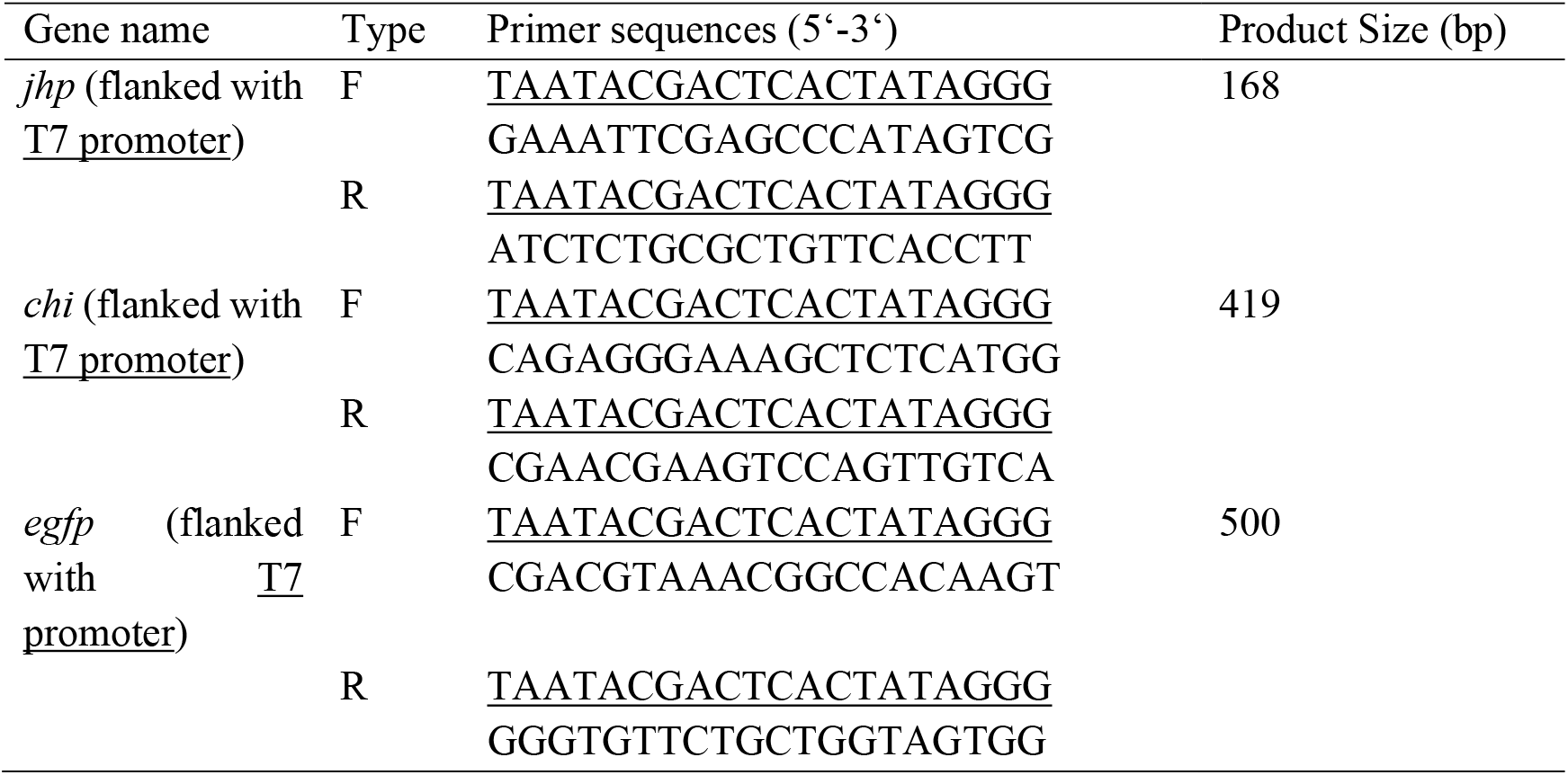
Primers designed for double-stranded RNA (dsRNA) synthesis.

### 2.5 Synthesis of layered double hydroxide (LDH) nanoparticles

LDH nanoparticles, were prepared using the co-precipitation and hydrothermal treatment method.^26^ Briefly, LDH particles were synthesized by vigorously mixing 10 mL of an aqueous solution containing magnesium chloride hexahydrate (MgCl_2_ * 6×H_2_O, Sigma-Aldrich, 99.0-102.0%, 3.0 mmol) and aluminum chloride hexahydrate (AlCl_3_ * 6H_2_O, Sigma-Aldrich, >97.0%, pellets) with 40 mL of sodium hydroxide solution (NaOH, Sigma-Aldrich, ≥97.0%, pellets) (0.15 M) for 10 min at room temperature. The LDH slurry was collected by centrifugation and then washed twice with double distilled water (40 mL), and resuspended in double distilled water (40 mL), after which the suspension was transferred to falcon tubes (15 mL) and hydrothermally treated at 100 °C for 16 h. The suspension contained approximately 1.4 mg × mL^-1^ of homogeneously dispersed LDH nanoparticles.

### 2.6 Electron microscopy

Copper grids of 400 mesh size, covered with pioloform and sputtered with carbon for stability were positioned directly or in dilutions (1:10, 1:100, 1:1000) for 5 min on a 20 µL drop of the suspension of nanoparticles described above. After three washing steps with ultrapure water to remove unbound material from the grid, the grids were air dried and analyzed at 80 kV in a Tecnai G2 Spirit (FEI, Frankfurt Germany, Thermo Fisher Scientific) transmission electron microscope equipped with a tungsten cathode. Images were taken with a side mounted CCD Veleta camera (Olympus, Münster, Germany) providing 2k × 2k pixel resolution. For image display and measurements served the TIA software (FEI). The 1:1000 dilution of nanoparticles suspension provided a suitable separation of hexagonal particles on the grids mesh for measurements. Obtained images were processed with Adobe photoshop CS4 extended, version 11, adjusting levels and brightness/contrast.

### 2.7 Double-stranded RNA loading on LDH

To determine the optimal and complete loading of respective dsRNA into LDH nanoparticles, the ratio of *in vitro* transcribed *jhp-, chi-* and *egfp-dsRNA* (500 ng) was combined multiple times with varying amounts of LDH (dsRNA: LDH from 1:2 to 1:9 [w/w]). dsRNA-LDH complex (BioClay^TM^) was loaded in a total volume of 10 μL and then incubated at room temperature for 30 min with gentle orbital agitation at 40 rpm. Complete loading of dsRNA onto LDH was confirmed by retention of the dsRNA-LDH complex in the well and unable to migrate through a 1% agarose gel at 80 V for 60 min in 1 × TAE buffer.^22,27^

### 2.8 RNase protection assay and pH stability test of the dsRNA-LDH nanoparticle complex

The ability of LDH to protect dsRNA was investigated by exposure of naked-dsRNA (500 ng) and the complex dsRNA-LDH (1:6) to RNase A (Ribonuclease A 90 U/mg [Kunitz], BioScience Grade, salt-free, Roth) treatment in three replicate experiments. Naked dsRNA and dsRNA-LDH were treated with different amounts of RNase A (0.15–0.000015 U) for 5 min at 37 °C. Then, dsRNA was released by acidic dissolution of LDH using release buffer (4.11 mL of 0.2 M Na_2_HPO_4_ + 15.89 mL 0.1 M citric acid; pH 3) for 10 min at a dsRNA-LDH buffer volume ratio of 5:1 and then loaded on a 1% agarose gel at 80 V for 60 min in 1 × TAE buffer.^22,27^ The initial RNase A amount of 0.15 U was diluted 10-, 100-, 1,000- and 10,000-fold which consisted of the five RNase enzyme amounts. Besides, the dsRNA-LDH complex was exposed to different pH levels (5, 9, and 11) at different time intervals (15, 30 and 60 min). pH stability experiment was conducted through gel electrophoresis using 0.1 M sodium acetate buffer (pH 5), 0.2 M sodium phosphate (pH 9), and 0.4 M sodium hydroxide (pH 11) and then loaded in total volume of 10 μL on a 1% agarose gel at 80 V for 60 min in 1 × TAE buffer.

### 2.9 *Phthorimaea absoluta* feeding bioassays

The effect of naked and formulated *jhp*- and *chi*-*dsRNA* was assessed against second instar *P. absoluta* larvae. A single tomato leaflet (4 mm long by 3 mm wide) excised with a razor blade from 6 weeks old tomato plant was placed in a Petri dish (60 mm diameter). Each leaflet was inoculated with either *jhp-dsRNA* or *chi-dsRNA* at a concentration of 500 ng × μL^-1^. Treated tomato leaflets were allowed to air dry at room temperature for 30 min and for each treatment, a single second instar *P. absoluta* larva was transferred to each leaflet using an artistic paintbrush. Each Petri dish represented a replicate and repeated three times per target gene. The Petri dishes were then sealed with Parafilm^®^ and transferred into plastic lunch boxes (17.5 × 12.5 × 6 cm) lined with moistened filter paper. Autoclaved distilled water and *egfp*-*dsRNA* were used as negative controls. Petri dishes were kept in an incubator at 25 ± 2 °C, 65 ± 5% relative humidity and 16 h light:8 h dark photoperiod. The treatments included feeding with: (1) autoclaved distilled water; (2) 3,000 ng of LDH; (3) 500 ng of *egfp*-dsRNA; (4) 500 ng of either naked *jhp-dsRNA* or naked *chi-dsRNA*; and (5) the complex of either *jhp-*dsRNA-LDH (BioClay *jhp-dsRNA*) or *chi*-dsRNA-LDH (BioClay *chi-dsRNA*) (1:6). All treatments were arranged in a completely randomized design with each treatment replicated three times as well as the controls. The experiment was performed twice under the same conditions using 10 second instar larvae (L2) per treatment, and three replicates were used for each treatment. Larvae were monitored daily for mortality for 10 days to observe the phenotypic effects, and the weight of the surviving pupae was measured with an analytical balance (Analysenwaage Kern^®^ 770, KERN & Sohn GmbH, Balingen-Frommern).

### 2.10 Real-Time quantitative PCR

To analyze the expression patterns of *jhp* and *chi*, total RNA was extracted from the *P. absoluta* larvae at 24, 48, and 72 h post feeding using the NEB Monarch Spin RNA extraction kit (New England Biolabs Inc., Frankfurt, Germany); each time point consisted of three biological replicates of 5 pooled larvae. Total RNA (1 µg) was reverse transcribed using the First Strand cDNA Synthesis Kit (Thermo Fisher Scientific, Waltham, MA, USA) according to the manufacturer’s instructions. First-strand cDNA synthesis was carried out using oligo (dT)_18_ primers. Real-time PCRs were carried out in a final volume of 10 μL, containing the Maxima SYBR Green qPCR Master Mix (2X) (5 μL) (Thermo Fisher Scientific, Germany), the primer pairs (500 nM) of each primer, the cDNA template (2 μL), and water (2 μL). Primers used in the analysis were validated with a standard curve based on a serial dilution of cDNA to determine the primer annealing efficiency. Reactions were carried out in three technical replicates in CFX96^TM^ Touch RT–PCR System (Bio-Rad, California, USA). For quantification of gene expression, the Pfaffl method^28^ was employed, using the *EF1*-α as reference gene and the control (water) larvae as calibrator for each time point. The amplification conditions were 95 °C for 10 min, followed by 36 cycles of 95 °C for 15 s, 56 °C for 30 s and 72 °C for 30 s.

### 2.11 Statistical analysis

Survival curves were analyzed using the Kaplan-Meier method, with differences assessed by the log-rank test. Pupal wieght data and gene expression between dsRNA-treated and control (water) were analyzed using one-way ANOVA, and the Tukey (HSD) test was used to compare means. Pupation rate and adult emergence data were analyzed with generalized linear model (GLM) using binomial distribution and logit link function. Time-mortality independent *t-*tests were performed to assess the effects of both formulated and naked dsRNAs on *chi* and *jhp* expression, pupal weight, and adult emergence rate, with the water group as the control. All statistical analyses were performed using R (version 3.6.3) statistical software packages^29^ and all statistical results were considered significant at the confidence interval of 95% (*P* < 0.05).

## 3 RESULTS

### 3.1 LDHs characterization and dsRNA loading to form dsRNA-LDH complex (BioClay^TM^)

Transmission Electron Microscopy (TEM) of the LDH crystals indicated a hexagonal nanosheet morphology, sizing of 60–170 nm (Figure 1A). Due to particle composition, thickness and density and associated electron scattering, optimal contrast was otained for imaging and required no additional staining with heavy metals to increase the contrast.^30^ The loading of *in vitro* transcribed dsRNA was performed in varying mass ratios of LDH nanoparticles. Successful loading was achieved by the absence of dsRNA migration from the wells during gel electrophoresis (Figure 1B). The complete loading of *jhp*-dsRNA on LDH nanoparticles at a mass ratio of 1:6 (Figure 1B) evidenced by the retention of all the dsRNA in the well compared with the control (Figure 1B, lane 2).

**Figure 1.**
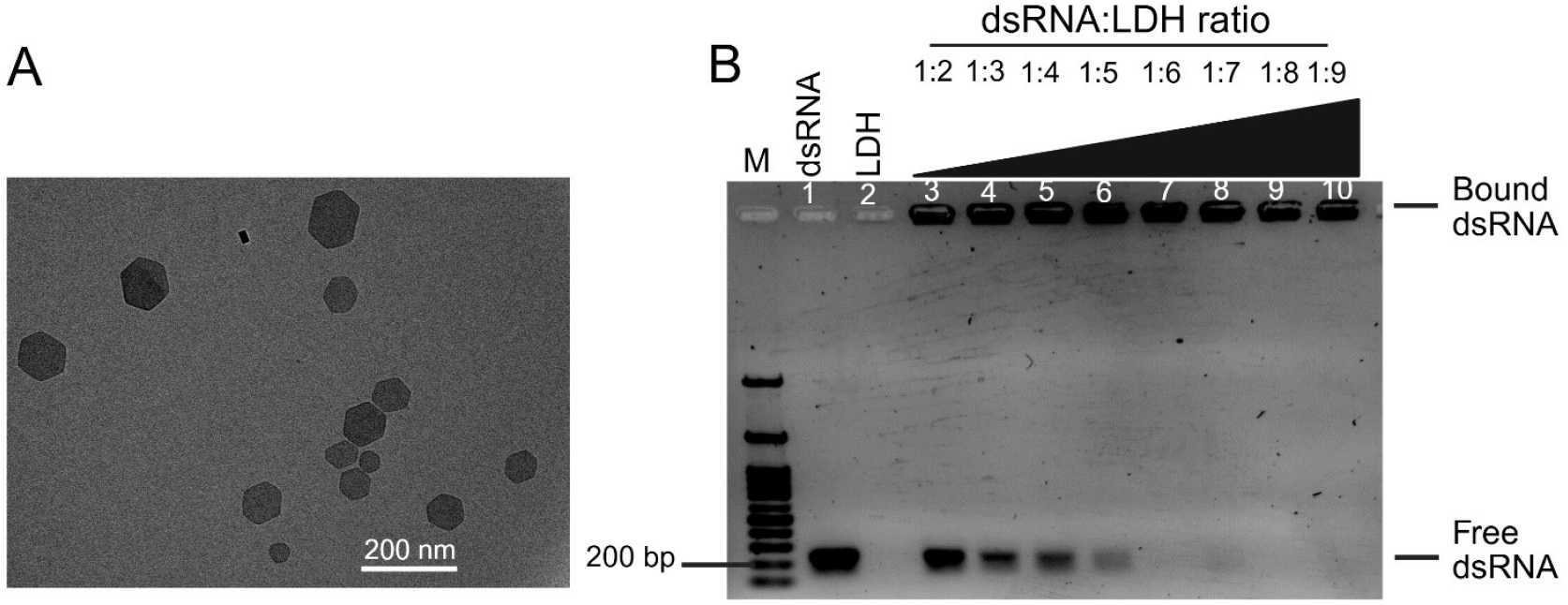
Characterization of hexagonal LDH nanoparticles and dsRNA loading into LDH. A) Transmission electron microscopy image of the 1:1000 diluted LDH suspension. The upper left projects the putative 3D shape. B) Loading profile of *jhp*-*dsRNA* onto LDH nanosheets. DsRNA was loaded into LDH nanoparticles using the mass ratios shown (dsRNA: LDH). Bioclay *jhp-dsRNA* preparations (1:2-1:9), 1 kb + DNA ladder (M), naked *jhp*-*dsRNA* (dsRNA), and LDH (LDH). LDH-bound dsRNA does not migrate and can be seen as fluorescence in the well while free dsRNA migrates through the agarose gel. Complete loading was achieved at a dsRNA:LDH mass ratio of 1:6.

### 3.2 RNase protection assay and pH stability test of the dsRNA-LDH nanoparticle complex

The capability of LDH nanoparticles to protect dsRNA from RNase degradation was tested through five different amounts of RNase A (0.15–0.000015 U). Higher RNase A amounts (0.15 U and 0.015 U) degraded the dsRNA either when naked (Figure 2, lanes 3 and 4) or complexed with the LDH nanoparticle compared to the control (Figure 2, lane 1). Conversely, lower RNase A amounts (0.0015 U–0.000015 U) did not degrade the dsRNA irrespective of the treatments (with or without LDH) (Figure 2, lanes 5-7 and lanes 10-12, for naked and formulated dsRNA, respectively).

**Figure 2.**
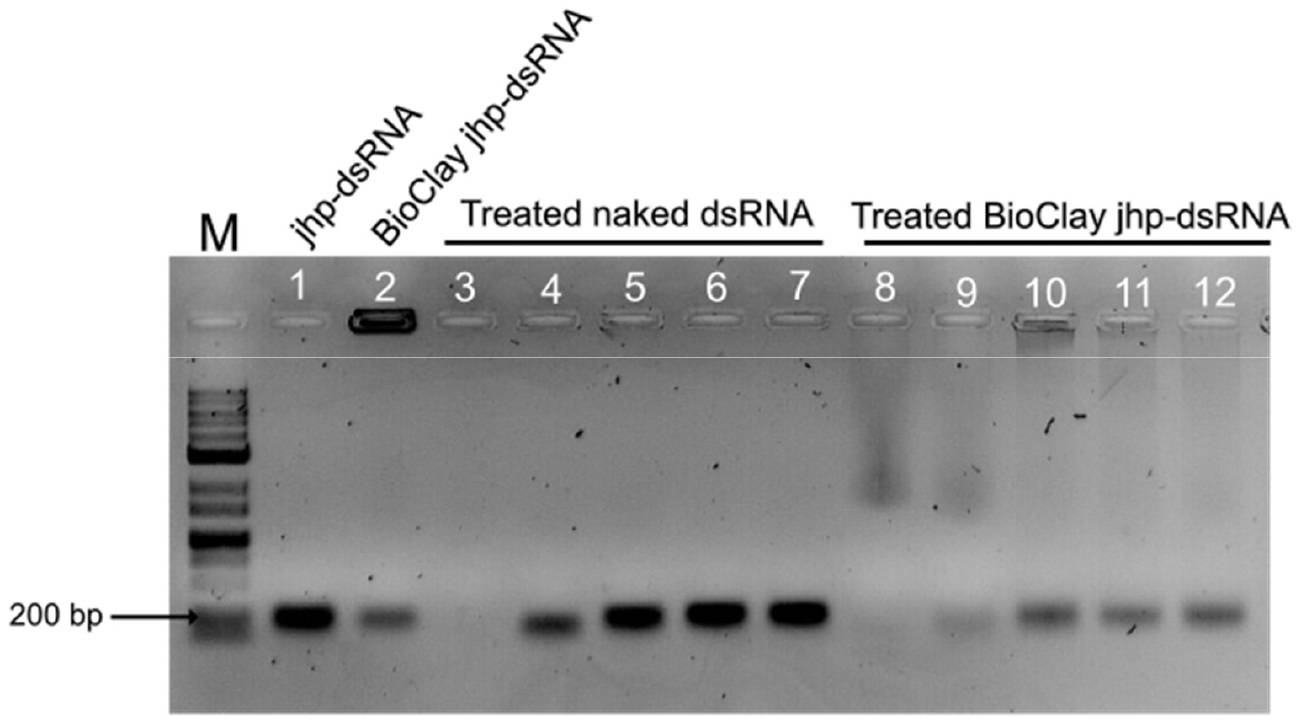
Treatment of naked dsRNA and dsRNA-LDH (Bioclay) complex with RNAse. The dsRNA of the treated and untreated dsRNA-LDH was released from LDH prior to gel electrophoresis. M = Marker (3 kb DNA ladder), Lane 1 = *jhp*-*dsRNA*, Lane 2 = BioClay^TM^ *jhp-dsRNA*, Lanes 3−7, naked dsRNA and lanes 8−12, dsRNA-LDH complex treated with decreasing RNase A amounts of 0.15 U, 0.015 U, 0.0015 U, 0.00015 U, and 0.000015 U.

The time-dependent pH stability assay of naked and dsRNA-LDH complex showed that dsRNA was not degraded at pH 5 irrespective of the incubation time (Figure 3). It is worth noting that the degradation of naked dsRNA was a little bit more pronounced at pH 11 compared to the control. For example, after 60 min of incubation, the band intensity of naked dsRNA was very weak compared to the control (Figure 3). Interestingly, part of dsRNA was released from the dsRNA-LDH complex at pH 9, and as pH increased from 9 to 11, the release of dsRNA was consistent (Figure 3).

**Figure 3.**
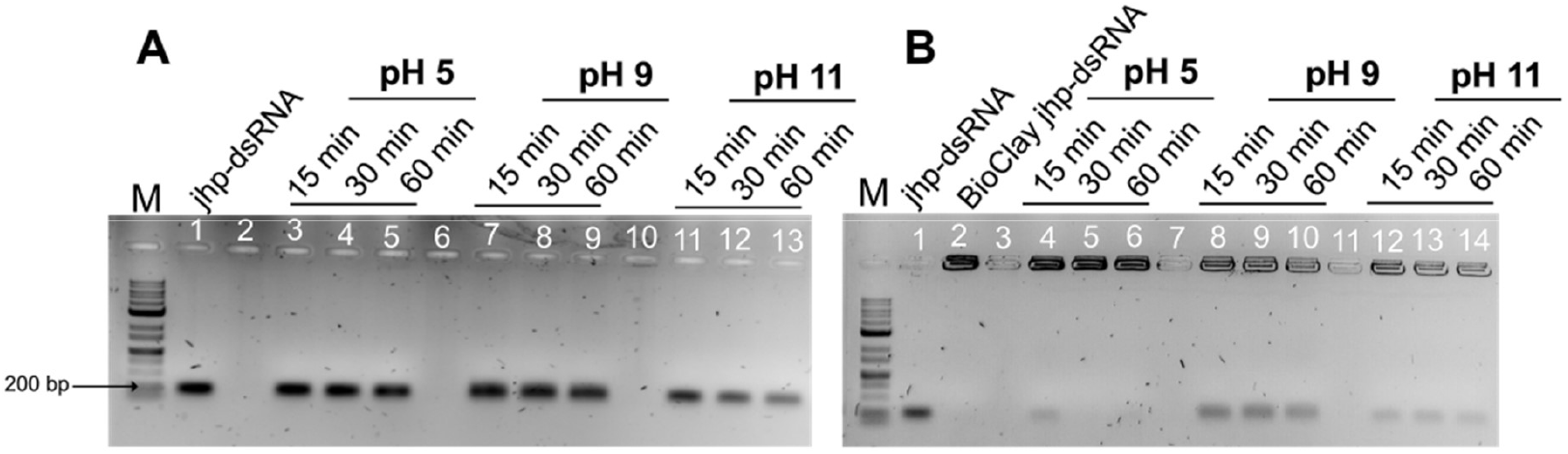
Gel electrophoresis showing release of dsRNA from LDH nanoparticles at varying pH and time intervals, mimicking insect gut conditions. (A) naked *jhp*-dsRNA incubated at different pH levels and times. (B) BioClay^TM^ *jhp*-*dsRNA* incubated at different pH levels and times. M = Marker (3 kb DNA ladder).

### 3.3 Survival of dsRNA-fed *Phthorimaea absoluta* larvae

Following tomato leaflet inoculation, no lethal effects on *P. absoluta* larvae of both formulated and naked dsRNA were observed (Log rank test = 10.5, df = 6, *P* = 0.14; Figure 4).

**Figure 4.**
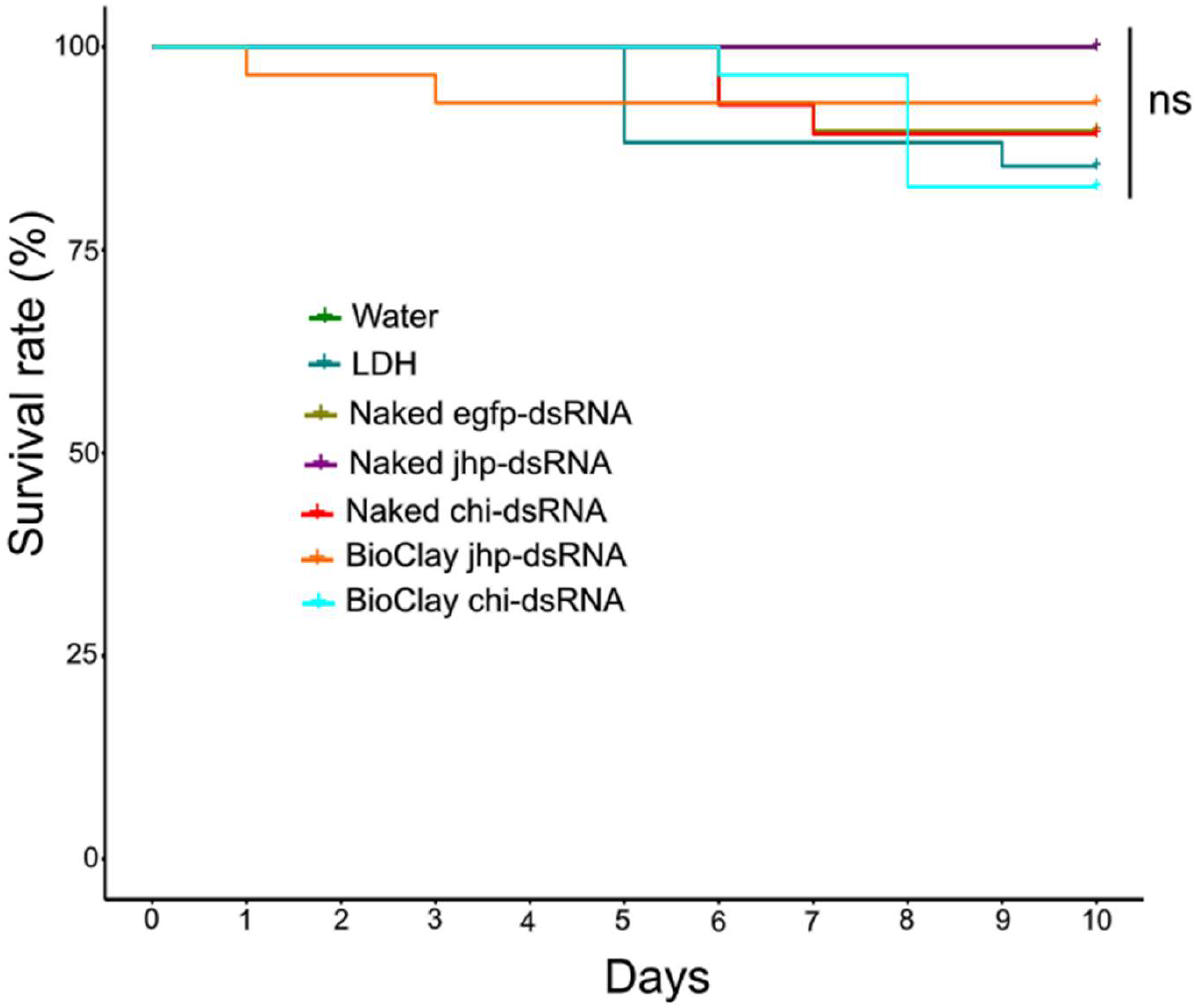
Survival analysis of *Phthorimaea absoluta* larvae across the different treatments. *jhp-dsRNA (juvenile hormone inducible protein)*; *chi-dsRNA* (*chitin synthase* A); *egfp-dsRNA (enhanced green fluorescent protein*); MgAl-LDH: Magnesium aluminum-layered double hydroxide nanoparticles. Each treatment group for the survival rate experiment had 30 individuals. Survival curves were analyzed using the Kaplan-Meier method, with differences assessed by the log-rank test. ns: no significant difference.

### 3.4 Silencing *chi* and *jhp* showed significant effects on *Phthorimaea absoluta* pupation rate, pupal weight, and adult emergence rate

No significant difference in pupation rate was observed among the treatments (χ^2^ = 0.73, df = 6, *P* = 0.99) (Figure 5 A). In contrast, significant differences in pupal weight were observed among the treatments compared to the control (F_6, 163_ = 2.96, *P* = 0.0091) (Figure 5B). The highest weight (4.29 ± 0.19 mg) was recorded on water samples while the lowest weight (3.42 ± 0.15 mg) was observed from larvae that fed on naked *jhp-dsRNA* followed by BioClay *jhp-dsRNA* (3.54 ± 0.15 mg) and BioClay *chi-dsRNA* (3.58 ± 0.14 mg) (Figure 5B). Upon emergence, significant difference in adult emergence rate was observed among the treatments compared to the control (water) (χ^2^ = 41.79, df = 6, *P* < 0.001) (Figure 6A). The highest emergence rate (96.29%) was recorded in the control (water) while naked *jhp-dsRNA* had the lowest adult emergence rate (39.94%) (t = -3.176, *P* = 0.0308) followed by BioClay *jhp-dsRNA* (42.61%) (t = -2.949, *P* = 0.0467) (Figure 6A and Figure 6B).

**Figure 5.**
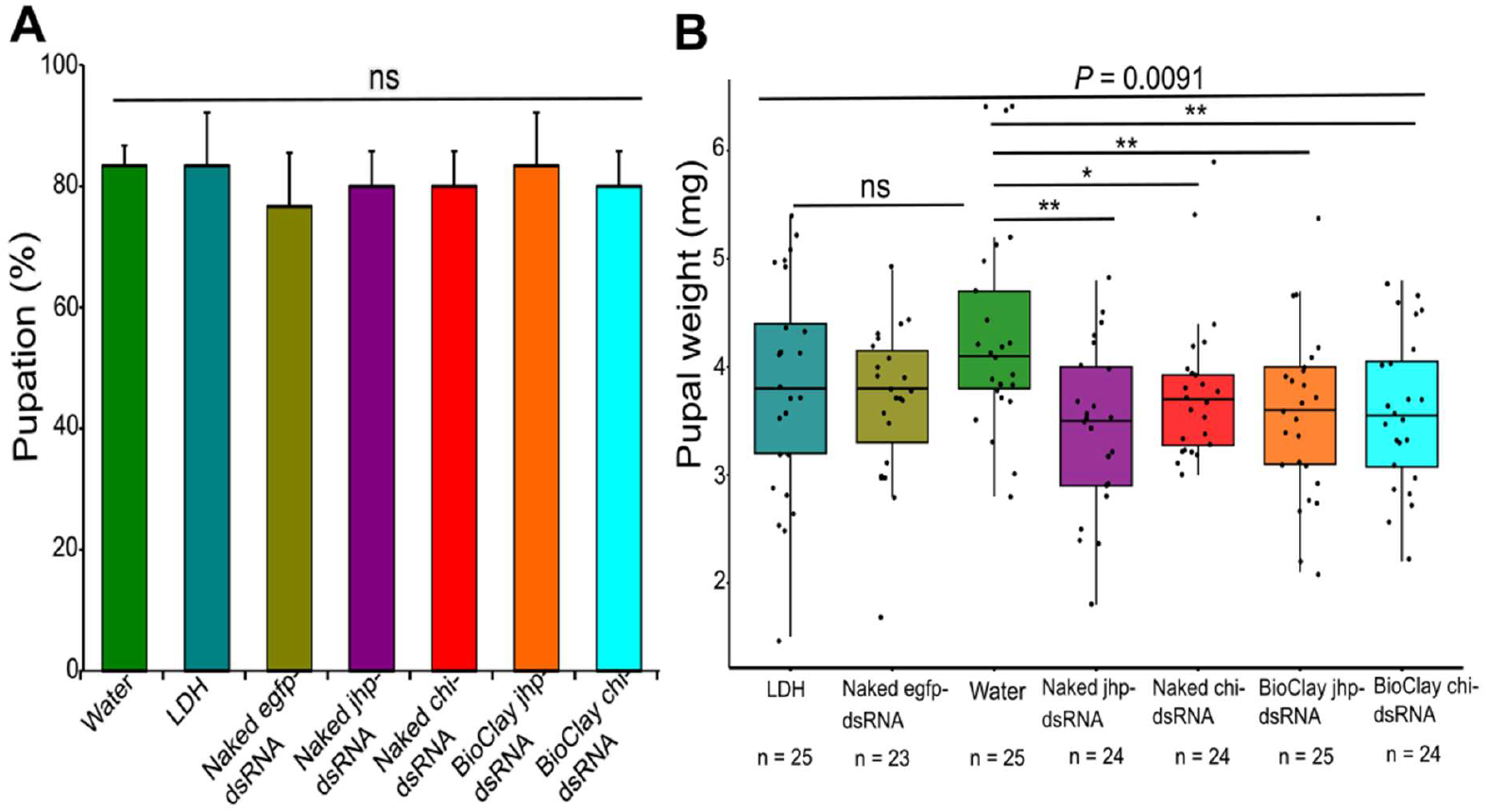
Effect of naked and BioClay^TM^ dsRNAs on *Phthorimaea absoluta* pupation and pupal weight (mg). A. Larvae that reached the pupal stage. B. Pupal weight. Asterisks denote the level of significance: * *P <* 0.05, ** *P <* 0.01 (one-way ANOVA followed by Tukey’s multiple comparisons). *ns*. no significant difference.

**Figure 6.**
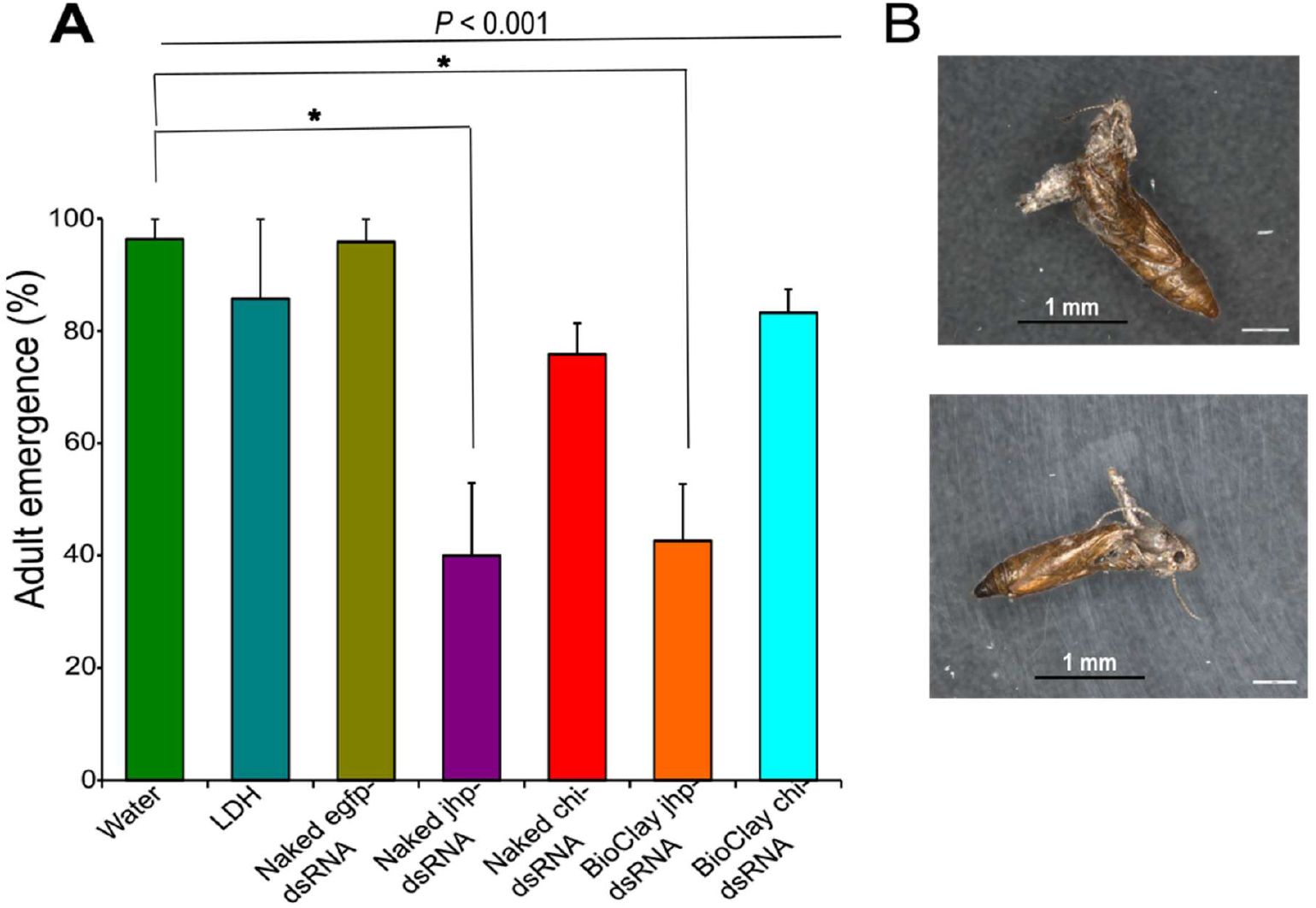
Effect of naked and BioClay^TM^ dsRNAs on *Phthorimaea absoluta* adult emergence. A. Adult emergence. B. Phenotypes of failed *P. absoluta* adults. Asterisks denote the level of significance: * *P <* 0.05, ** *P <* 0.01 (one-way ANOVA followed by Tukey’s multiple comparisons).

### 3.5 Double-stranded RNA-LDH complex significantly reduced expression of the target genes in *P. absoluta*

Relative expression levels of *jhp* and *chi* mRNAs were determined at 24, 48, and 72 h. Both formulated and naked dsRNA were effective at suppressing *chi* and *jhp* expression. For example, at 24 h post-feeding, there was 53 and 57% reduction in transcript abundance of *chi* gene in larvae that fed either on naked *chi-dsRNA* or BioClay *chi-dsRNA*, respectively, compared with the control (Figure 7A). In constrast, after 48 h, larvae fed on naked *chi-dsRNA* did not show any significant reduction in mRNA levels while a 71% reduction seen in the BioClay *chi-dsRNA* treatment compared to the control (water) (Figure 7B). Interestingly, a significant reduction of 73% and 48% was observed in accumulation of *chi* transcripts in larvae fed with *chi-dsRNA* and BioClay *chi-dsRNA* after 72 h, respectively (Figure 7C).

**Figure 7.**
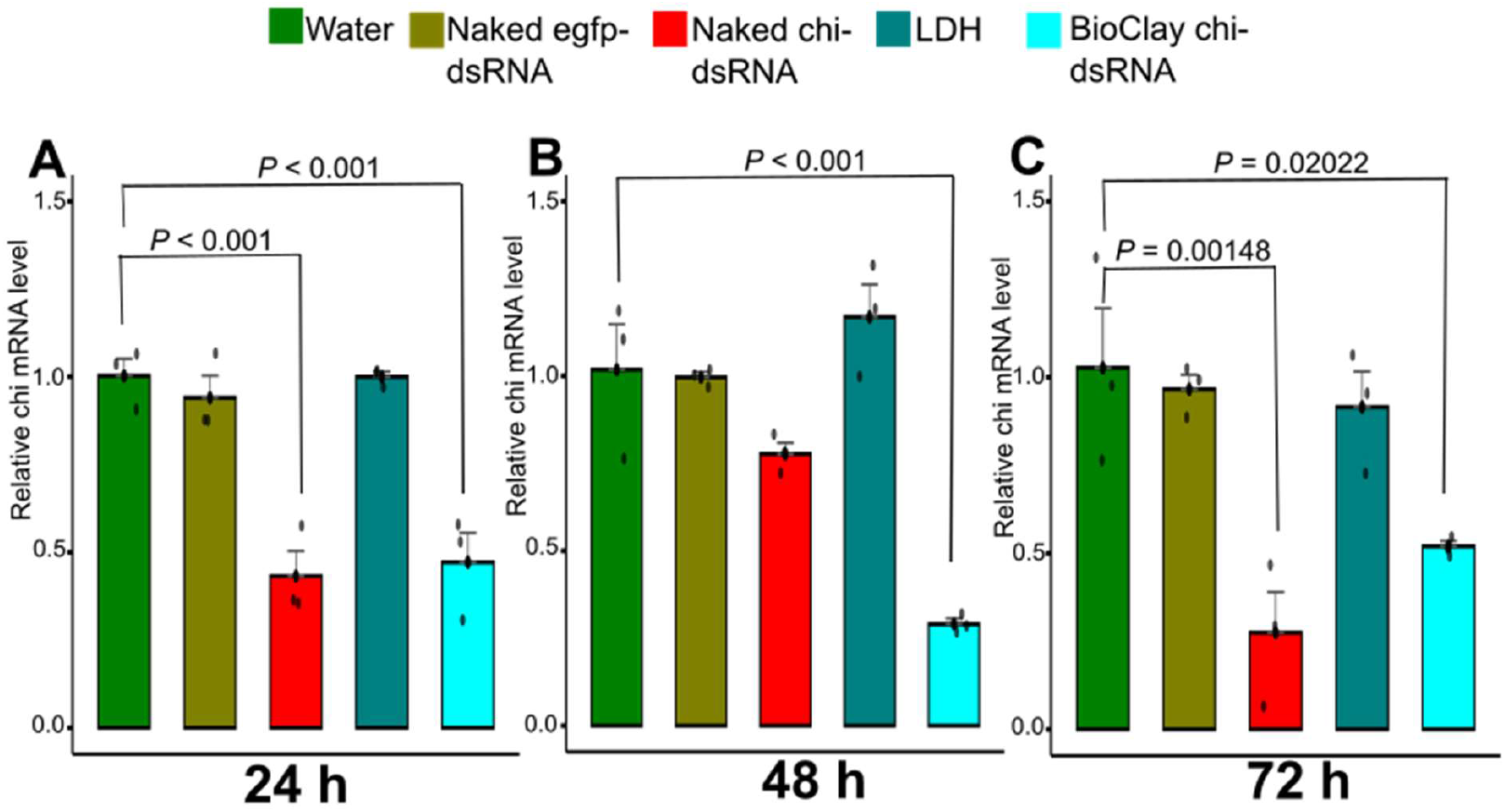
*Chi* gene mRNA level in *Phthorimaea absoluta*. Expression levels normalized to *EF1-α* gene. A) after 24 h post-feeding, B) after 48 h, and C) after 72 h. Error bars represent standard error of the mean of 3 technical replicates (5 insects per replicate). Each black dot represents a single data point (observation). Significant differences between means (multiple t-tests) are depicted.

The expression levels of *jhp* were significantly reduced by 35 and 46% in *P. absoluta* larvae fed with *jhp*-*dsRNA* and BioClay *jhp-dsRNA* after 24 h, respectively (Figure 8A). Conversely, no significant reduction in *jhp* mRNA levels was observed in larvae fed with BioClay *jhp-dsRNA* while a 45% reduction was recorded in the naked *jhp-dsRNA* compared to the control after 48 h of active feeding (Figure 8B). At 72 h post-feeding, a 39% reduction in transcript abundance of *jhp* gene was observed only in larvae fed on BioClay *jhp-dsRNA* (Figure 8C).

**Figure 8.**
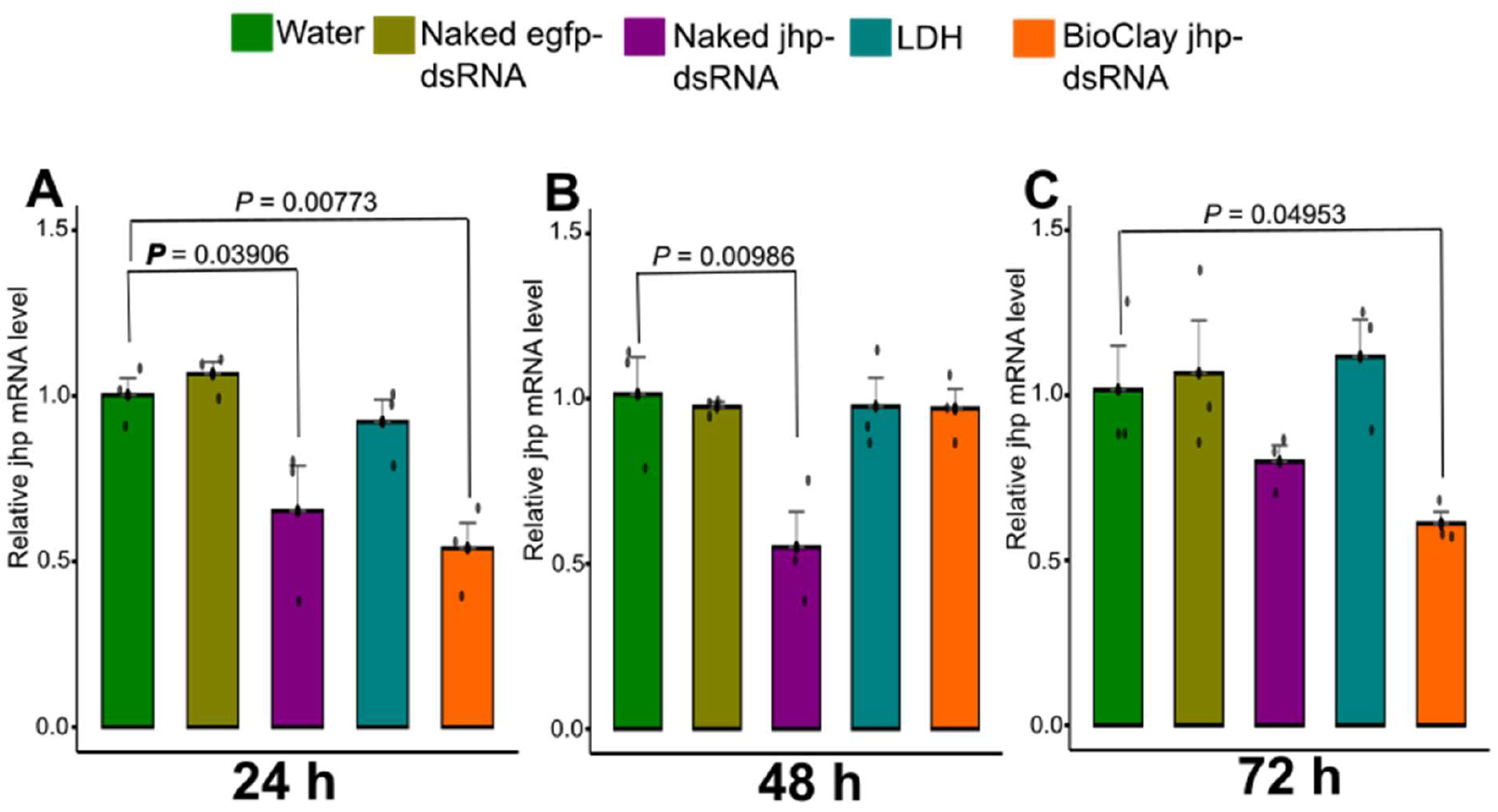
*Jhp* gene mRNA level in *Phthorimaea absoluta*. Expression levels normalized to *EF1*-*α* gene. A) after 24 h post-feeding, B) after 48 h, and C) after 72 h. Error bars represent standard error of the mean of 3 technical replicates (5 insects per replicate). Each black dot represents a single data point (observation). Significant differences between means (multiple t-tests) are depicted.

## 4 DISCUSSION

Our study demonstrates that MgAl-LDH nanoparticles can adsorb dsRNA to form dsRNA– LDH complexes, referred to as BioClay^TM^, providing to some extent stability to dsRNA and protection against nuclease activity. We also showed that formulated and naked dsRNAs targeting *jhp* and *chi* genes did not cause significant impact on the survival and pupation rate of *P. absoluta*, but triggered important sublethal phenotype changes through the reduction of pupal weight and adult emergence. The findings indicate that the use of RNAi technology against these target genes yielded developmental/physiological impairment rather than acute lethality, highlighting the importance of deploying RNAi-based control agents as a component of a potential strategy for *P. absoluta* management.

MgAl-LDH nanoparticles stand out as one of the nanocarriers possessing excellent physical properties to deliver dsRNA to target insect pests.^31^ Here, we loaded dsRNA onto Mg Al-LDH nanoparticles and achieved complete loading when the mass ratio of dsRNA/LDH was 1:6. Then, when performing dsRNA degradation activity, we could show that higher RNase A amounts degraded the dsRNA either when naked or complexed with the LDH nanoparticles while lower RNase A amounts did not degrade the dsRNA irrespective of the treatments. Further, our findings revealed that dsRNA is stable over a distinct time period when in buffer solutions at pH 5 and pH 9 at room temperature. As the pH increased from 9 to 11, which represents the alkaline environment in *P. absoluta* midgut, we observed partial release of dsRNA from the nanoparticles while naked dsRNA showed slight degradation. This indicates that dsRNA can be released from the dsRNA/LDH complex in alkaline environments as a result of chemical hydrolysis. Indeed, the stability of RNA toward chemical and enzymatic degradation is of fundamental importance for a successful uptake of the dsRNA by the target insects.^32^ Under alkaline pH conditions, we expected the dsRNA-LDH complex to resist chemical hydrolysis, which is a key factor for successful delivery of dsRNA to *P. absoluta*. However, the release of dsRNA from the nanoparticles may indicate that alkaline pH conditions trigger the availability of free RNA ready for uptake by the cells in *P. absoluta*. This is consistent with the observations by Jiang *et al*.^33^ who developed a pH-responsive nanoparticle, chitosan-polyethylene glycol-carboxyl (CS-PEG-COOH), capable of releasing dsRNA under alkaline conditions, thereby enhancing the delivery of dsRNA against larvae of beet armyworm, *Spodoptera exigua* (Hübner) (Lep.: Noctuidae). It is worth mentioning, that the most attractive and important property of LDHs is that their interlayer anions are exchangeable, indicating that various organic anions, inorganic anions, and nucleic acids can be intercalated into the interlayer of LDHs to attribute different functions.^34^

We found that following tomato leaflet inoculation, formulated and naked dsRNAs targeting *jhp* and *chi* genes did not cause rapid larval mortality compared to the controls (water, LDH, and naked eGFP). In contrast, Askew *et al*.^21^ demonstrated significant decrease in larval mortality when delivering Ryanodine receptors (RyRs) dsRNA complexed with chitosan nanoparticles to *P. absoluta* following 10 days feeding compared to naked *TaRy* dsRNA. Also, Bento *et al*.^19^, using bacterially-expressed dsRNA, demonstrated considerable larval mortality in *P. absoluta* when targeting *jhp* and *chi* genes. Similarly, when delivering naked *V-ATPase-A* (*vacuolar atpase-a*) and *Arginine kinase* (*ak*) dsRNAs at a tested dose of 5,000 ng to *P. absoluta* larvae feeding on tomato leaflets, Camargo *et al*.^17^ showed significant larval mortality and reduction of leaf damage. Yet, except for Bento *et al*.^19^, these authors used higher concentrations of dsRNA ranging from 3 to 5 µg/µL to achieve lethal effect in *P. absoluta*. Clearly, using lower concentrations of dsRNA, as in our study, might not always be as effective. We therefore recommend that future studies explore the use of different doses and specific concentrations of dsRNA.

Interestingly, our results showed that even though both formulated and naked dsRNA did not cause significantly affect survival of *P. absoluta* larvae and pupation rate, we noted, however, a significant effect of dsRNA on pupal weight and adult emergence. This was evidenced by lighter pupae in the case of either formulated or naked *chi-* and *jhp-dsRNA*, resulting in malformed pupae or adults, which were ultimately found dead. The reduced pupal weight indicates that dsRNA was successfully taken up and ultimately triggered gene silencing in *P. absoluta*. Interestingly, even using naked dsRNA (without a carrier) appears to have some biological activity, suggesting that *P. absoluta* has some susceptibility to RNAi. Majidiani *et al*.^20^ also reported significant reduction in *P. absoluta* pupal weight when targeting acetylcholinesterase (*AChE*), nicotinic acetylcholine alpha 6 (*nAChRs*), and ryanodine (*RyRs*) receptors through injection and root delivery of dsRNAs. Similarly, Li *et al*.^35^a showed significant reduction in adult emergence when *P. absoluta* larvae fed on tomato leaflets inoculated with dsRNA that target two essential genes including the *voltage-gated sodium channel* (*Nav*) and *NADPH-cytochrome P450 reductase* (*CPR*). Conversely, Bento *et al*.^19^ did not detect significant differences in pupal weight between treatments, even for those dsRNAs that led to high larval mortality and significant reduction in the accumulation of transcripts of the target genes including *jhp* and *chi*. It is worth noting that JH is a key endocrine regulator that is essential for the growth, metamorphosis and reproduction in insects.^36,37^ On the other hand, chitin synthases are large membrane proteins known to catalyze the polymerization of N-acetylglucosamine into chitin.^38^ Chitin is a major component of the exoskeletons and peritrophic membranes of insects. Consequently, down-regulation of *jhp* and *chi* expression might lead to the defects in the pupal-adult transition that we observed in our present study (Figure 6B). However, the proportion of dead individuals was not statistically different for larvae that fed on *chi*-dsRNA compared to the control. This is surprising as the gene expression pattern over time revealed a significant reduction in transcript abundance of the *chi* gene in larvae that fed on either naked dsRNA or BioClay compared to the control. Hence, a successful RNAi outcome is probably more determined by the nature of the target gene than its silencing efficiency, underlining the importance of target gene selection. Similar results were obtained by Chen *et al*.^38^ who observed that even though injection of *chitin synthase A* dsRNA into the larvae of *S. exigua* resulted in developmental arrest and cuticle and trachea defects, some larvae managed to have new cuticles, and chitin in the cuticle suggesting the hypothesis that the RNAi was not completely effective. Further studies are necessary to explore whether these phenotypes are the result of a higher RNAi efficacy and to elucidate where exactly this silencing is happening.

Our study showed that oral devlivery of dsRNA via feeding mediated by clay nanoparticles resulted in variable silencing of *jhp* and *chi* genes. Measurement of target gene expression at various time points after dsRNA delivery revealed a significant decrease as early as 24 h for both *chi* and *jhp* genes delivered either naked or in a formulated way. Conversely, after 48 h, larvae that fed on either naked *chi-dsRNA* or BioClay *jhp-dsRNA* did not show any significant reduction in mRNA levels. This pattern probably reflects the transient effect of the dsRNA in the cells as a result of its lower initial uptake. On the other hand, one would expect that using clay nanosheets as carrier to deliver dsRNA to *P. absoluta* may induce more lethal effect.

Clearly, besides the alkaline pH conditions that may influence the dsRNA stability, as demonstrated in this study, another important factor that affects RNAi efficiency in lepidoptera is the presence of double-stranded ribonuclease (dsRNase) in the digestive tract, which can degrade dsRNA prior to uptake by gut cells.^39^ For example, Li *et al*.^40^ recently identified in *S. exigua* four double-stranded ribonucleases (SedsRNase1, SedsRNase2, SedsRNase3, and SedsRNase4) whose inhibition significantly decrease dsRNA degradation ability, contributing to high efficiency of RNAi. We observed after 72 h a significant reduction of 73% and 48% in accumulation of *chi* transcripts in *P. absoluta* larvae fed with *chi-dsRNA* and BioClay *chi-dsRNA*, respectively. One potential explanation is that gene silencing only occurs in a particular section of the gut.^41^ Hence, further studies are needed to elucidate the uptake mechanism of the dsRNA-LDH complex into *P. absoluta* midgut cells.

In conclusion, our study demonstrates that using dsRNA-based control agents to target *jhp* and *chi* candidate genes in *P. absoluta* can be feasible. However, improved delivery methods will need to be developed to improve RNAi efficiency. A long-term sustainable management strategy of *P. absoluta* should entail the promotion of different compatible and environmentally-friendly control methods.

## ACKNOWLEDGEMENTS

The first author would like to thank the Alexander von Humboldt Foundation for supporting the research. Authors are also grateful to the University of Bonn and the Julius Kühn Institute for additional financial support. We thank Verena Maiberg for her excellent technical assistance in electron microscopy. The graphical abstract was created in BioRender. Agbessenou, A. (2025) https://BioRender.com/c3vt2nf.

## DATA AVAILABILITY STATEMENT

Research data are included in the article.

